# Temperature Sensitive Glutamate Gating of AMPA-subtype iGluRs

**DOI:** 10.1101/2024.09.05.611422

**Authors:** Anish Kumar Mondal, Elisa Carrillo, Vasanthi Jayaraman, Edward C. Twomey

**Author notes:** Correspondence (E.C.T.).

## Abstract

Ionotropic glutamate receptors (iGluRs) are tetrameric ligand-gated ion channels that mediate the majority of excitatory neurotransmission^1^. iGluRs are gated by glutamate, where upon glutamate binding, they open their ion channels to enable cation influx into post-synaptic neurons, initiating signal transduction^2^. The structural mechanism of iGluR gating by glutamate has been extensively studied in the context of positive allosteric modulators (PAMs)^3–15^. A fundamental question has remained – are the PAM activated states of iGluRs representative of glutamate gating in the absence of PAMs? Here, using the α-amino-3-hydroxy-5-methyl-4-isoxazolepropionic acid subtype iGluR (AMPAR) we show that glutamate gating is unique from gating in the presence of PAMs. We demonstrate that glutamate gating is temperature sensitive, and through temperature-resolved cryo-electron microscopy (cryo-EM), capture all major glutamate gating states. Physiological temperatures augment channel activation and conductance. Activation by glutamate initiates ion channel opening that involves all ion channel helices hinging away from the pores axis in a motif that is conserved across all iGluRs. Desensitization occurs when the local dimer pairs decouple and enables closure of the ion channel below through restoring the channel hinges and refolding the channel gate. Our findings define how glutamate gates iGluRs, provide foundations for therapeutic design, and point to iGluR gating being temperature sensitive.

## Introduction

Neuronal cells communicate with each other at synapses via neurotransmitters. The major neurotransmitter in the brain is glutamate ^1,16^. Glutamate is released from pre-synaptic neurons and received by ionotropic glutamate receptors (iGluRs) in the post-synaptic neuronal membrane. iGluRs are tetrameric ligand-gated ion channels and upon binding glutamate open their ion channels to enable cation entry into the post-synaptic neuron^1,2^. This initiates depolarization and is the cornerstone of excitatory neurotransmission. Thus, iGluRs are critical for healthy brain function, and their dysregulation is linked to many neurological disorders including Alzheimer’s, Parkinson’s, epilepsy, chronic pain, schizophrenia, ataxia, and glioblastoma^1,17–19^.

There are four major iGluR subtypes in humans. These include the α-amino-3-hydroxy-5-methyl-4-isoxazolepropionic acid (AMPA), N-methyl-D-aspartate (NMDA), kainate, and delta subtypes. All subtypes share a common architecture: an extracellular domain (ECD) comprised of an amino terminal domain (ATD) and ligand-binding domain (LBD), where glutamate binds, and below the ECD, a transmembrane domain (TMD) that houses an ion channel^1,2^. Below the TMD is a carboxy terminal cytosolic domain that enables membrane localization at synapses. The chief role of the ATD is for tetrameric assembly and interaction with cross-synaptic adhesion factors^1^. The LBD is clamshell-shaped, and each half of the LBD (D1 and D2) close around glutamate upon binding. In general, it is this action that opens the ion channel below via direct linkers between the LBD and TMD^2^.

AMPA-subtype iGluRs (AMPARs) are the fastest iGluR subtype. Upon binding glutamate, they allow cation influx on the single millisecond timescale^2^. This fast depolarization is critical for rapid information processing in the brain^16^. AMPAR signaling is tuned throughout the brain by different combinations of AMPAR subunits GluA1-4 in the core tetrameric subunit, and transmembrane AMPAR regulatory proteins (TARPs) that affect AMPAR gating kinetics and synaptic localization^1,16,20^.

Decades of electrophysiology and structural studies have formed our current understanding of AMPAR gating^1,2^. AMPARs have three main functional states: resting, open, and desensitized. Glutamate binding to the AMPAR LBD initiates the gating cycle, where AMPARs enter a primed state – the LBD clamshells are closed around the neurotransmitter, but the conformational changes that accommodate activation or desensitization have not yet occurred. Activation of the AMPAR occurs when the lower part of the LBD, D2, moves up to the LBD upper half, D1, to fully close around glutamate. When this occurs in local LBD dimers of the AMPAR tetramer, the D1’s of LBDs contact each other, creating a D1-D1 interface. This coordinated motion pulls on the LBD-TMD linkers to open the ion channel. The receptor desensitizes to protect the cell from excitotoxicity when the LBD clamshells are maximally closed around glutamate, but instead of pulling on the TMD, the D1-D1 interface is decoupled, and an interface between LBD D2’s is formed. This D2-D2 interface relaxes tension on the LBD-TMD linkers, putting the ion channel in a closed state despite glutamate being fully bound.

There is a caveat to this gating paradigm. No activated AMPARs have been captured in the presence of only glutamate, nor for any iGluR^3–15^. But what if the glutamate activation mechanism is unique from the activation mechanism in the presence of PAMs? This is a major missing piece of information. To this end, we sought to capture a bona fide activated state of an AMPAR in the presence of only glutamate.

We hypothesized that temperature may play a role in augmenting glutamate gating of AMPARs. It has long-been appreciated that temperature augments excitatory post-synaptic potentials from cephalopods through mammals^21–23^. We thought AMPAR gating may be directly temperature sensitive. Pointing to this are early studies showing that temperature increases AMPAR ligand affinity^24,25^ and the suggestion that acceleration of AMPAR gating kinetics may play a role in altering post-synaptic currents during temperature increases^26^.

To this end, we definitively demonstrate that AMPAR gating is temperature sensitive, and we captured each major AMPAR gating state using temperature resolved cryo-EM. The glutamate-activated state is distinct from the PAM-activated states of AMPARs. During glutamate activation, an activation seal between LBDs holds open the ion channel pore, where all four ion channel helices hinge away from the pore axis. Hinging of the ion channel helices occurs in a two-fold symmetric manner in a motif conserved across all iGluRs. Desensitization occurs when the LBD activation seal is ruptured, which relaxes the pull on the ion channel helices, closing the pore. Our results provide new foundations for understanding glutamate gating of iGluRs, drug design, and point to iGluR gating being temperature sensitive, which is an important consideration when considering hyperthermic temperatures in traumatic brain injury and neurological disorders^27–29^.

### Physiological temperatures augment AMPAR gating

To test the idea that AMPAR gating is temperature sensitive, we used the GluA2-γ2 construct, which mimics synaptic AMPAR function^3,6,7,30,31^. GluA2-γ2 is a fusion construct where TARPγ2, an AMPAR auxiliary subunit that promotes channel opening, is fused via its N-terminus to the carboxy-terminus of GluA2 (modified rat GluA2_flip_, edited to Q at the Q/R site, Methods).

We performed temperature steps (25 °C to 30 °C to 37 °C to 42 °C) and recorded currents from single channels in the presence of 10 mM glutamate (Fig. 1a, Table 1, Methods). These data generally shows that unitary channel currents activated by glutamate are more frequent and larger when temperature is increased (Fig 1a). These observations are GluA2-γ2 dependent (Extended Data Fig. 1). AMPARs conduct to multiple sub-conductance levels, where GluA2-γ2 complexes show four sub-conductance levels (O1-O4)^6,32^. Time course fittings of the channel openings reveal how occupancy of O1-O4 is temperature dependent (Fig. 1b, Table 1). The sub-conductance levels we observe are O1 (∼12 pS), O2 (∼35 pS), O3 (∼45 pS), and O4 (∼65 pS). At room temperature (25 °C), the current histogram is best fitted with two Gaussian components, corresponding to two conductance levels, the mean amplitudes (and populations) of which are 12 ± 1.7 pS (5 ± 0.4 %) and 35 ± 5.2 pS (90 ± 5 %). At 30 °C, the current histogram and Gaussian fitting reveals three conductance levels: 12 ± 1.1 pS (4.5 ± 0.5 %), 35 ± 0.9 pS (8 ± 0.6 %), and 45 ± 4.8 pS (84 ± 4.5 %). Stepping to healthy physiological temperature (37 °C) further augments channel sub-conductance, where four conductance levels are observed: 12 ± 1.3 pS (4.5 ± 0.6 %), 35 ± 3.7 pS (10 ± 1.2 %), 45 ± 5.1 pS (14 ± 1.6 %), and 65 ± 2.4 pS (62 ± 6.1 %). Channels at hyperthermic temperature (42 °C) have increased occupancies of the higher conductance levels, which is exemplified by 72 ± 7 % occupancy of the highest sub-conductance level, O4. The overall changes in channel sub-conductance are reflected in the mean unitary channel conductance, where the mean conductance is ∼40 pS at 25 °C, ∼50 pS at 30 °C, ∼60 pS at 37 °C, and ∼70 pS at 42 °C (Fig. 1c).

**Fig. 1.**
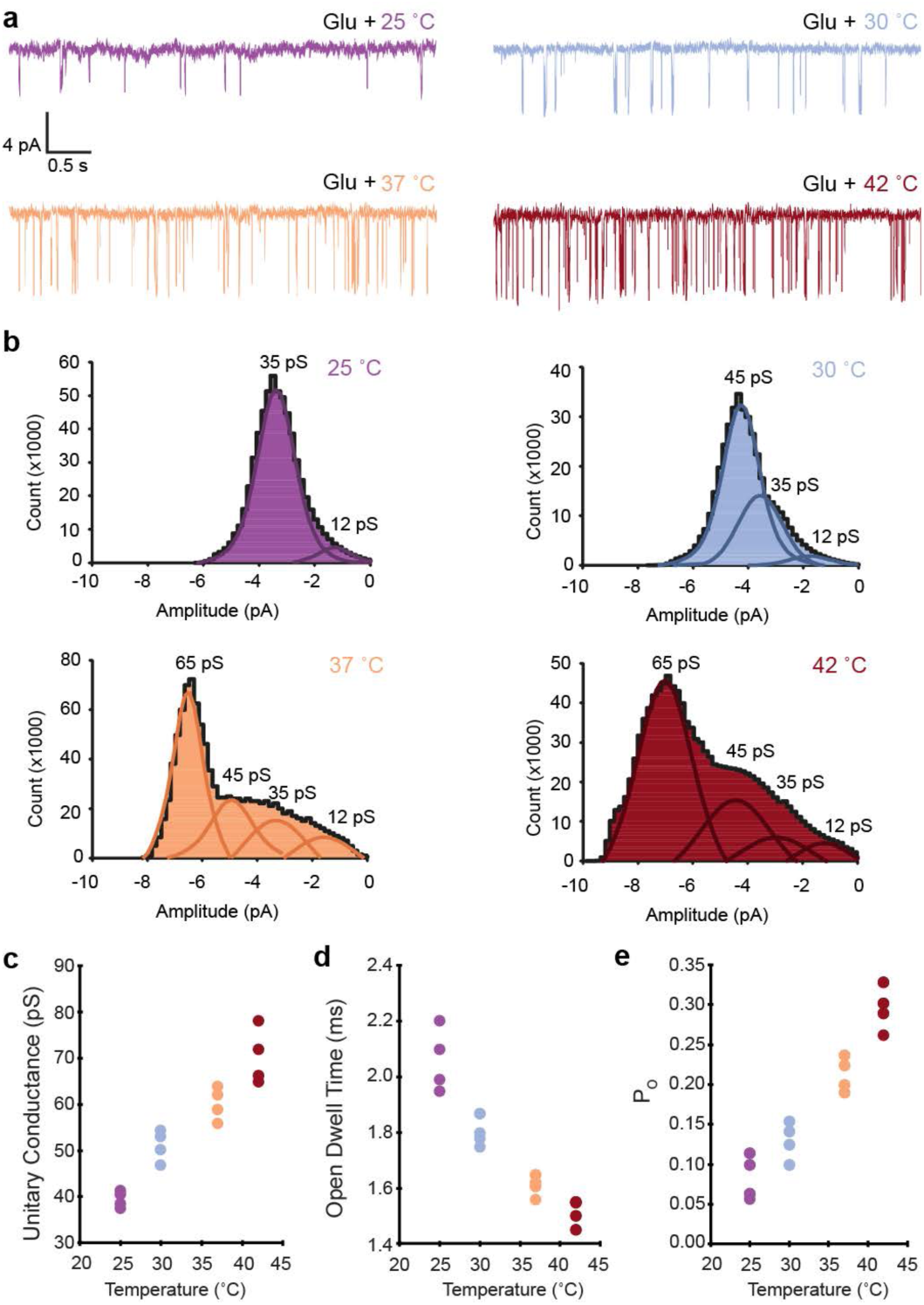
Temperature augments AMPAR function. **a**, Example GluA2-γ2 single channel currents at 25 °C, 30 °C, 37 °C, and 42 °C recorded on a single membrane patch in the presence of 10 mM glutamate (Glu). **b**, Current histograms with Gaussian fits for the amplitude events. Four conductance levels (12 pS, 35 pS, 45 pS, 65 pS) were observed. **c**, Average unitary conductance, **d** average open dwell time, **e** channel open probability (P_O_) for single channel membrane patches at each temperature (n=4 patch current recordings for each 25 °C, 30 °C, 37 °C, 42 °C). The amplitude events were recorded for 1-2 min at each temperature step per patch. The total times recorded for each temperature step were: 8 min (25 °C), 5 min (30 °C), 7 min (37 °C), and 5 min (42 °C).

**Table 1.** Summary statistics of GluA2-γ2 single channel recordings. Percentage of occurrence for each level or event is shown next to its value in parentheses. The error in the lifetimes is the SE of the fit. The table is related to Fig 1.

| Conductance level (pS) | 25 °C | 30 °C | 37 °C | 42 °C |
| --- | --- | --- | --- | --- |
| 1 | 12 ± 1.7<br>(5 ± 0.4 %) | 12 ± 1.1<br>(4.5 ± 0.5 %) | 12 ± 1.3<br>(4.5 ± 0.6 %) | 12 ± 0.6<br>(6 ± 0.7 %) |
| 2 | 35 ± 5.2<br>(90 ± 5 %) | 35 ± 0.9<br>(8 ± 0.6 %) | 35 ± 3.7<br>(10 ± 1.2 %) | 35 ± 1.3<br>(9 ± 1.3 %) |
| 3 | --- | 45 ± 4.8<br>(84 ± 4.5 %) | 45 ± 5.1<br>(14 ± 1.6 %) | 45 ± 2.7<br>(12 ± 1.1 %) |
| 4 | --- | --- | 65 ± 2.4<br>(62 ± 6.1 %) | 65 ± 2.4<br>(72 ± 7 %) |
| **Mean open time (ms)** | 2.06 ± 0.05 | 1.8 ± 0.02 | 1.6 ± 0.02 | 1.5 ± 0.02 |
| **Open events** |  |  |  |  |
| τ1 | 1.2 ± 0.2<br>(78 ± 5 %) | 0.8 ± 0.06<br>(65 ± 4.6 %) | 0.65 ± 0.04<br>(60 ± 5 %) | 0.3 ± 0.05<br>(64 ± 3.3 %) |
| τ2 | 4.5 ± 0.3<br>(15.5 ± 2 %) | 3.9 ± 0.5<br>(30 ± 3.1 %) | 2.9 ± 0.7<br>(32.5 ± 2.2 %) | 2.5 ± 0.2<br>(27 ± 5.5 %) |
| **Mean Shut time (ms)** | 138.2 ± 27 | 106 ± 18 | 75.6 ± 10.5 | 34 ± 6 |
| **Shut events** |  |  |  |  |
| τ1 | 17.5 ± 3.7<br>(29 ± 7 %) | 13.9 ± 0.85<br>(18 ± 3 %) | 8.2 ± 1.2<br>(33 ± 1.6 %) | 3.7 ± 0.8<br>(41 ± 2.9 %) |
| τ2 | 39 ± 5<br>(23 ± 5 %) | 22.5 ± 3.5<br>(35 ± 5 %) | 16.9 ± 3.3<br>(40 ± 5.4 %) | 10.7 ± 2.6<br>(39 ± 4.3 %) |
| τ3 | 382 ± 62<br>(26 ± 8%) | 292 ± 35<br>(26 ± 8.5 %) | 206 ± 24<br>(17 ± 2.4 %) | 173 ± 11.5<br>(10 ± 2.2 %) |

The 25 °C unitary conductance is in line with previous single channel recordings performed on single AMPAR-TARP complexes at room temperature^6,33–35^. Physiological temperature (37 °C) markedly increases the average conductance, with hyperthermic temperature (42 °C) showing a ∼1.75x increase in unitary conductance compared to room temperature. However, inversely, the open channel dwell time decreases as temperature increases from room temperature to hyperthermic by approximately 0.5 ms (Fig. 1d), as does the mean shut time of the channel (Table 1). Juxtaposed to this, there is a dramatic change in channel open probability (P_O_) as temperature increases (Fig. 1e). At room temperature, the open probability is approximately 8.4%. At physiological temperature, this raises to 21.3%, and at hypothermic temperature, the P_O_ raises to 29.6%.

In total, this data suggests that AMPAR gating is augmented by increased temperatures. While the relative open dwell time is decreased, unitary channel conductance increases, as does P_O_. The increase in channel conductance and decrease in mean open time are consistent with the increase in ion mobility and reaction rates upon increasing temperature as predicted by the Arrhenius equation. This is consistent with temperature-dependent gain of function observed in excitatory post-synaptic currents^36^ and the premise that temperature increases state transitions and ion movements in ion channels^26,37–40^.

On the other hand, increase in P_O_ is indicative of a shift in equilibrium between the closed and open states of the receptor. This shift reflects a change in the Gibbs free energy (ΔG) between the closed and open states, with an increase of the fraction of receptors in the open channel state at higher temperatures. Our observations are consistent with the prediction that temperature may drive AMPARs into higher sub-conductance levels^26^.

Based on our direct evidence, we conclude that AMPAR function is temperature sensitive, and that temperature should be a critical consideration in electrophysiological and drug design studies against AMPARs.

### Temperature resolved cryo-EM

Because temperature positively influences the gating of GluA2-γ2, we hypothesized that raising the temperature of GluA2-γ2 during cryo-EM specimen preparation would enable reconstruction of the rare glutamate-activated state. We refer to raising temperature during cryo-EM specimen preparation to capture short-lived biological states, that are temperature sensitive and rarely occurring during the typical specimen preparation temperatures (e.g., 4 °C or room temperature) as temperature-resolved cryo-EM. Indeed, temperature-resolved cryo-EM has been utilized to resolve the temperature gating of TRP channels^41–44^ and temperature-dependent enzymatic ensembles^45^. We adopted a similar approach here (Methods).

Because hyperthermic conditions increase GluA2-γ2 activation compared to 37 °C, we prepared GluA2-γ2 for temperature-resolved cryo-EM at 42 °C. We purified GluA2-γ2 and tested the thermostability of GluA2-γ2, we analyzed the complex stability using fluorescence-based size exclusion chromatography (FSEC) after incubating GluA2-γ2 for fifteen minutes at 4 °C, 37 °C, and 42 °C (Extended Data Fig. 2). There was no major difference in the stability of GluA2-γ2 between the temperature incubations as observed by maintenance of the GluA2-γ2 complex peak. We prepared GluA2-γ2 for cryo-EM using hyperthermic conditions (Methods). In brief, GluA2-γ2 was primed for glutamate gating by first incubating the specimen at 42 °C on a heat block (Fig. 2a). Immediately prior to vitrification, GluA2-γ2 was spiked with 1 mM glutamate (pre-heated to 42 °C) and vitrified using a cryoplunger with the specimen chamber set to 42 °C. Parallel to this, we prepared GluA2-γ2_EM_ at 42 °C without glutamate to capture the apo or resting state of the receptor (Fig. 2a). The micrographs from GluA2-γ2 prepared at hyperthermic conditions enable definitive identification of single particles (Fig. 2b), and distinct views as revealed by two-dimensional (2D) classification (Fig. 2c).

**Fig. 2.**
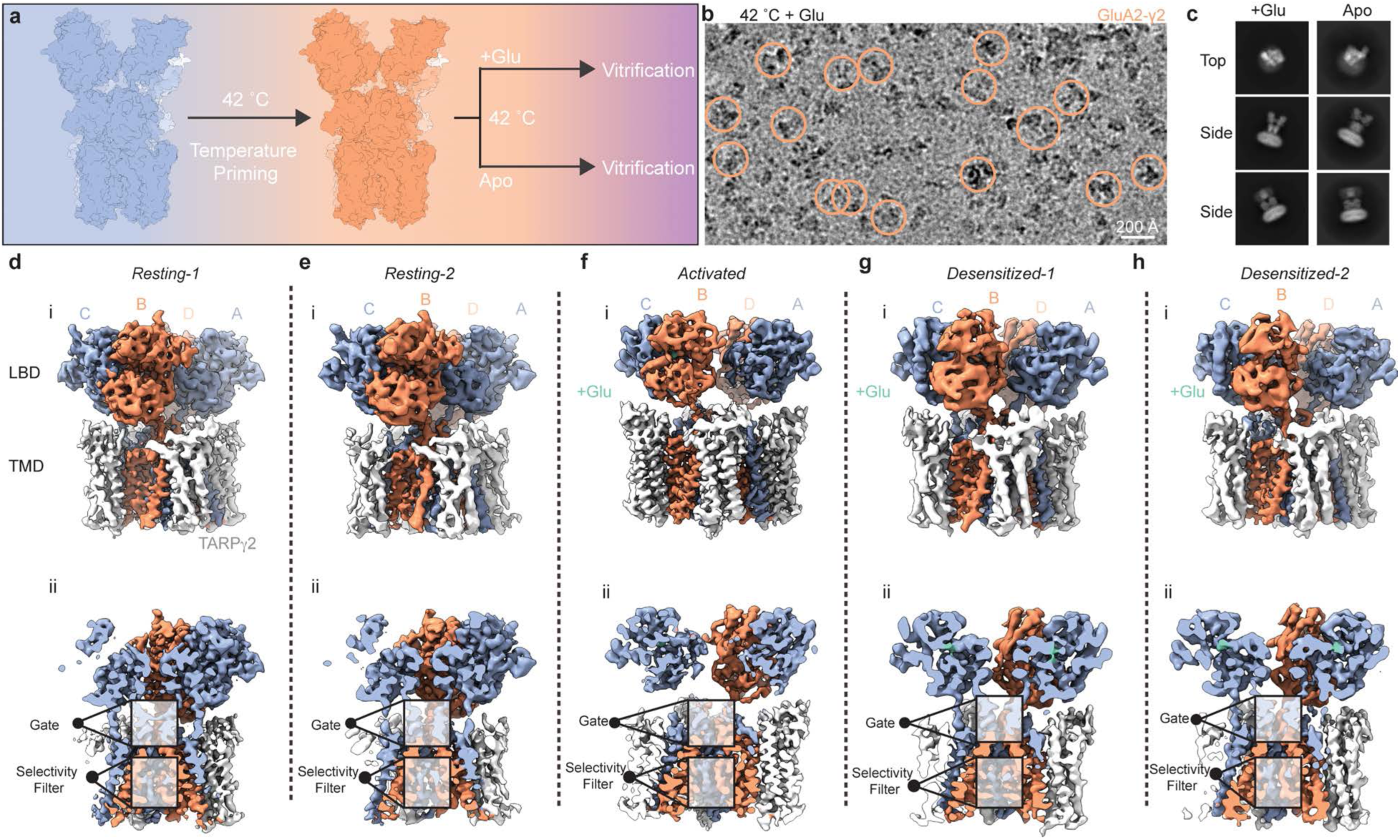
Temperature-resolved cryo-EM. **a**, Workflow for temperature-resolved cryo-EM. **b**, Example micrograph of AMPARs vitrified after hyperthermic temperature priming and 1 mM Glu exposure. **c**, Example 2D classes from temperature-resolved cryo-EM in the presence and absence of Glu. **d-h**, Ensemble of states reconstructed from temperature-resolved cryo-EM. For each panel d-h: inset i, full view of cryo-EM map; inset ii, map sliced to central pore axis.

To parse out the distinct conformational states, we used signal subtraction to remove the AMPAR ATD, focused on the AMPAR LBD and TMD in three-dimensional (3D) classifications and refinements (Methods, Extended Data Figs. 3-6, Table 2). This revealed a gating ensemble (Fig. 2d-h), with two resting states (resting-1, resting-2), an activated state, and two desensitized states (desensitized-1, desensitized-2). The resting-1 LBD-TMD was refined to 4.18 Å, with the central TMD reaching 3 Å locally (Extended Data Fig.6a,b). The resting-2 LBD-TMD was refined to 4.78 Å, with the central TMD reaching 3.5 Å locally (Extended Data Fig.6a,b). The activated state LBD-TMD was refined to 3.54 Å, and TMD to 3.46 Å, with the central TMD reaching ∼2.5 Å locally (Extended Data Fig. 4a,b). The desensitized-1 LBD-TMD was refined to 4.48 Å, TMD to 4.43 Å, reaching ∼3.5 Å locally in the central TMD (Extended Data Fig. 4a,b). The desensitized-2 LBD-TMD was refined to 4.31 Å, and the TMD to 4.02 Å, with the central TMD approaching ∼3.0 Å locally (Extended Data Fig. 4a,b).

**Table 2:** Cryo-EM data collection, refinement, and validation statistics.

|  | Resting-1<br>(EMDB-46872)<br>(PDB 9DHP) | Resting-2<br>(EMDB-46873)<br>(PDB 9DHQ) | Activated<br>(EMDB-46874)<br>(PDB 9DHR) | Desensitized-1<br>(EMDB-46875)<br>(PDB 9DHS) | Desensitized-2<br>(EMDB-46876)<br>(PDB 9DHT) |
| --- | --- | --- | --- | --- | --- |
| **Data collection and processing** |  |  |  |  |  |
| Magnification | 130,000x | 130,000x | 130,000x | 130,000x | 130,000x |
| Voltage (kV) | 300 | 300 | 300 | 300 | 300 |
| Electron exposure (e<sup>-</sup>/Å<sup>2</sup>) | 40 | 40 | 40 | 40 | 40 |
| Defocus range (μm) | -1.0 – 2.5 | -1.0 – 2.5 | -1.0 – 2.5 | -1.0 – 2.5 | -1.0 – 2.5 |
| Pixel size (Å) | 0.97 | 0.97 | 0.97 | 0.97 | 0.97 |
| Symmetry imposed | C2 | C2 | C2 | C2 | C2 |
| Initial particle images (no.) | 1,524,514 | 1,524,514 | 3,320,119 | 3,320,119 | 3,320,119 |
| Final particle images (no.) | 74,785 | 38,242 | 56,570 | 44,872 | 43,184 |
| Map resolution (Å) | 4.18 | 4.78 | 3.54 | 4.48 | 4.31 |
| FSC = 0.143 |  |  |  |  |  |
| Map resolution range (Å) | 2.6 – 15.0 | 2.9 – 12.3 | 2.3 – 11.0 | 2.8 – 12.9 | 2.5 – 12.1 |
| **Refinement** |  |  |  |  |  |
| Initial model used (PDB code) | 5WEO | 5WEO | 5WEO | 5WEO | 5WEO |
| Model resolution (Å) | 4.3 | 4.8 | 3.7 | 4.5 | 4.3 |
| FSC = 0.143 |  |  |  |  |  |
| Model resolution range (Å) | 3.8 – 4.3 | 3.8 – 4.7 | 3.4 – 3.7 | 4.2 – 4.5 | 3.9 – 4.4 |
| Map sharpening *B* factor (Å<sup>2</sup>) | -137 | -151 | -94 | -150 | -134 |
| Model composition |  |  |  |  |  |
| Non-hydrogen atoms | 18075 | 18075 | 18304 | 18296 | 18296 |
| Protein residues | 2328 | 2328 | 2350 | 2350 | 2350 |
| Ligands |  |  |  |  |  |
| *B* factors (Å<sup>2</sup>) |  |  |  |  |  |
| Protein | 9.79/157.56/70.15 | 33.9/254.14/135.00 | 5.32/246.11/135.00 | 126.66/328.61/229.51 | 12.51/196.38/94.31 |
| Ligand |  |  |  |  |  |
| R.m.s. deviations |  |  |  |  |  |
| Bond lengths (Å) | 0.003 | 0.002 | 0.014 | 0.003 | 0.003 |
| Bond angles (°) | 0.670 | 0.714 | 0.837 | 0.714 | 0.665 |
| Validation |  |  |  |  |  |
| MolProbity score | 1.52 | 1.55 | 1.45 | 1.53 | 1.47 |
| Clashscore | 3.23 | 3.17 | 3.1 | 3.11 | 2.94 |
| Poor rotamers (%) | 0.16 | 0 | 0.72 | 0.2 | 0.31 |
| Ramachandran plot |  |  |  |  |  |
| Favored (%) | 93.98 | 93.01 | 95.03 | 93.52 | 94.30 |
| Allowed (%) | 5.84 | 6.51 | 4.62 | 5.79 | 5.35 |
| Disallowed (%) | 0.18 | 0.48 | 0.35 | 0.69 | 0.35 |

Each state has four TARPγ2 molecules bound around the AMPAR TMD, as expected. The state of each LBD can be identified, along with AMPAR subunit position, transmembrane (TM) helix, ion channel gate, and selectivity filter.

The differences between the cryo-EM maps represent different functional states of each class. Glutamate is not bound to the AMPAR LBDs in the resting states, which have a closed channel gate (Fig. 2d,e). In the activated state, glutamate is bound to the AMPAR LBDs, and the channel gate is open (Fig. 2f). In the desensitized states, glutamate is bound, but the channel gate is closed (Fig. 2g,h).

Thus, temperature-resolved cryo-EM enabled us to capture the major gating states of GluA2-γ2 in the presence or absence of only glutamate.

### Glutamate activation mechanism

In the activated state, all four LBDs are bound to glutamate, and four molecules of TARPγ2 are bound (Fig. 3a). The receptor is overall two-fold symmetric. The LBDs are organized into two local dimers labeled by subunit position (dimer pairs A & D, B & C), of which A/C and B/D are symmetric pairs.

**Fig. 3.**
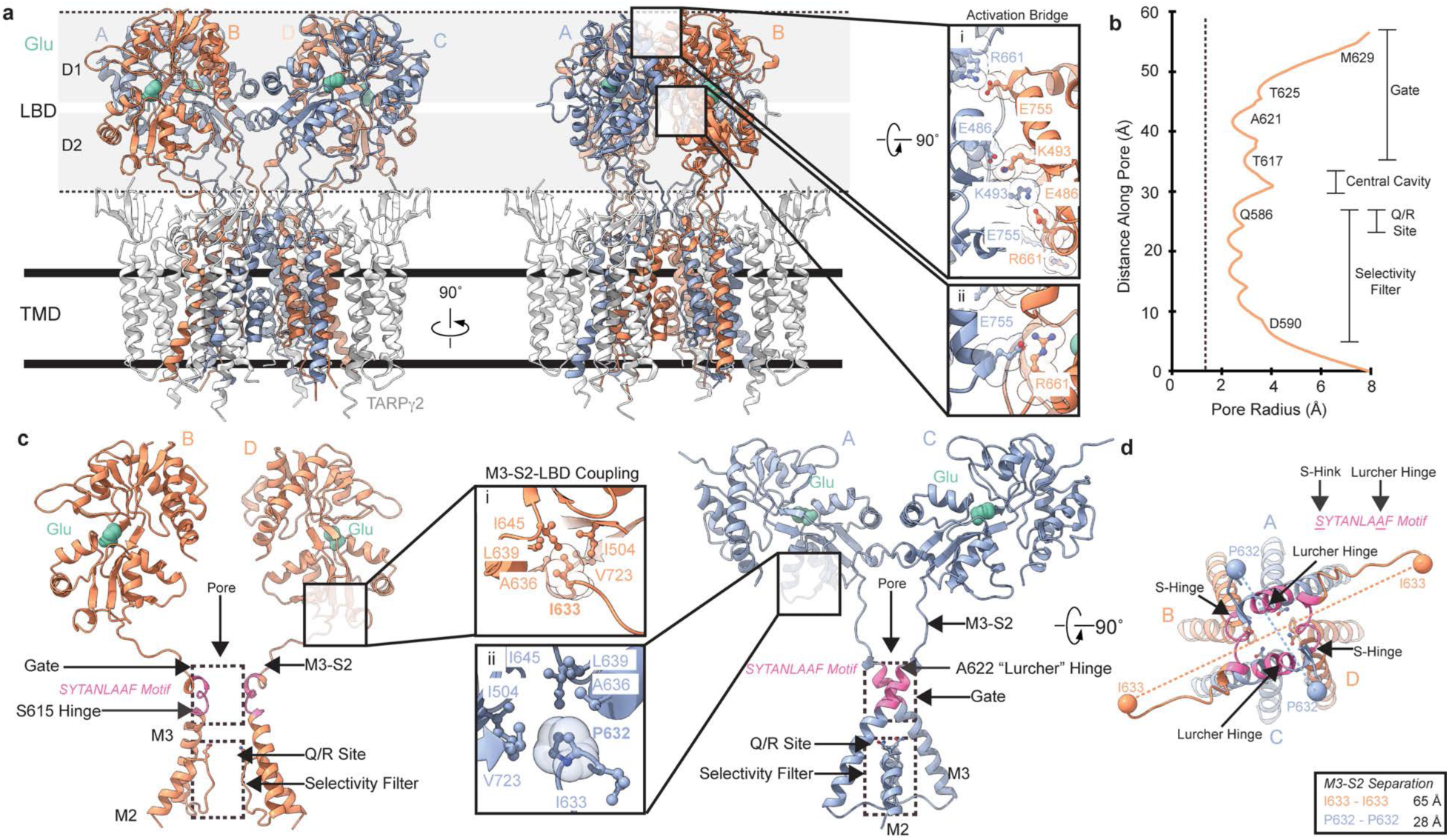
Glutamate activation. **a**, Structure of the GluA2-γ2 glutamate-activated state. Insets i, ii are close ups of the salt bridges that collectively make the activation bridge. **b**, pore radius profile (orange) along the ion channel pore. Residues are marked immediately next to the plot, with general features on the right. Dashed line represents the 1.4 Å radius of a water molecule. **c**, Close up of the symmetric subunit pairs in the glutamate-activated state. B/D subunits, left; A/C subunits, right. Inset i is a zoom on the M3-S2-LBD coupling in B/D, inset ii in A/C. **d**, top view of the glutamate-activated ion channel pore, rotated 90° from panel c with all four subunits.

There is marked contact between LBDs in their local dimers, where salt bridges between the LBD upper halves (D1) contact each other extensively (Fig. 3a, inset i). Here, the central salt bridges are E486 from (A/C) to K493 (B/D) and K493 (A/C) to E486 (B/D). On the far side of the central salt bridges is a contact between D2 and D1, where R661 (A/C) contacts E755 (B/D) (Fig. 3a, inset i). The symmetric pair of this interaction occurs on the outer side of the LBD dimer, where E755 (A/C) is in contact with R661 (B/D) (Fig. 3a, insets i & ii). Collectively, we refer to the group of four salt bridges as the activation bridge. And indeed, the activation bridge is essential for activation. Ablating the salt bridges within the activation bridge markedly reduces the ability of glutamate to hold the channel open, dramatically increasing AMPAR desensitization^46,47^.

During glutamate binding, the lower D2 half of an LBD swings upward to the D1 half to close around glutamate. Therefore, the R661-E755 salt bridges help to maintain the glutamate-bound state of each LBD by locking the D2 in the glutamate-bound position against its LBD partner.

The activation bridge holds apart the LBD D2’s within local LBD dimers. Collectively, this puts tension on the LBD-TMD linkers below (Fig. 3a), which opens the ion channel. As a result, the ion channel is in a putative open state (Fig. 3b). The ion channel gate, at the top of M3 helices, is formed by M629, T625, and T617 is pulled away from the central axis and does not occlude the pore. The minimal constructive radius (r_min_) at the gate is approximately 2.5 Å. Below, at the selectivity filter, the r_min_ is 2.3 Å. The channel radius at the Q/R site is 2.5 Å. Thus, the pore is expected to be in a conductive state for water and cations such as sodium and calcium.

The open ion channel is held open by the M3-S2 linkers which directly tether the ion channel gate to the LBDs. In the B and D subunit positions, the linkers are held in place by the linker residue I633 being locked into a hydrophobic cavity on the bottom of D2 (Fig. 3c, inset i). In the A/C subunit positions, M3-S2 LBD coupling occurs via P632 insertion into the hydrophobic cavity (Fig. 3c, inset ii). Collectively, this puts tension on the M3-S2 linkers that causes unwinding of each M3 helix in all subunits, opening the ion channel for conductance (Fig. 3d).

The features of the cryo-EM map allow clear identification of the features in the unwound pore helices (Extended Data Fig. 7b). Unwinding of the M3 helices is facilitated by a hinge at S615 in the B/D subunits and A622 in the A/C subunits. In GluA2-γ2 S615 is at the beginning of the SYTANLAAF motif, which is a conserved motif in mammalian iGluRs (Extended Data Fig. 8a). A622 is the penultimate residue in the motif. Interestingly, A622 is the site of the “Lurcher” mutation (A622T) that causes spontaneous firings in iGluRs and leads to excitotoxic cell death^48–52^, and has long been of great interest to the field since its discovery in 1997^48^. Perhaps the Lurcher mutation exacerbates the hinging of the subunits in the A/C subunit positions. Thus, we term the hinging at A622 in the A/C subunits the Lurcher hinge, and the hinging at S615 in the B/D subunits the S-hinge.

While hinging in all four M3 helices is unique to this glutamate-activated AMPAR structure, it has been proposed to perhaps be necessary for maximum conductance of the ion channel^6,12^. Furthermore, the two-fold symmetric opening of the channel via the S-Hinge and Lurcher hinge agrees with early biochemistry suggesting the that the SYTANLAAF motif imparts a two-fold symmetric gate in iGluR ion channels^53^.

The SYTANLAAF motif has long been proposed to be critical for iGluR gating^54,55^. Our data shows that the entire SYTANLAAF motif is ingrained in gating, and points to why mutations in this motif are so debilitating across the iGluR family^1,19,55^.

### Glutamate activation is distinct from PAM activation

To delineate whether glutamate gating is distinct from gating in the presence of PAMs, which all previous activated AMPAR states have been studied in the presence of, we compared our glutamate activated state to a putative PAM-activated state, GluA2-γ2 in the presence of CTZ and glutamate^3,6^ (pdb 5WEO).

Both the activated state and PAM-activated state are two-fold symmetric (Fig. 4a,b). In each, all four LBDs are bound to glutamate, but in the PAM-activated state, CTZ is wedged between LBDs (Fig. 4b) to lock in the activated state and prevent desensitization. The major differences between the states lay within the ion channel.

**Fig. 4.**
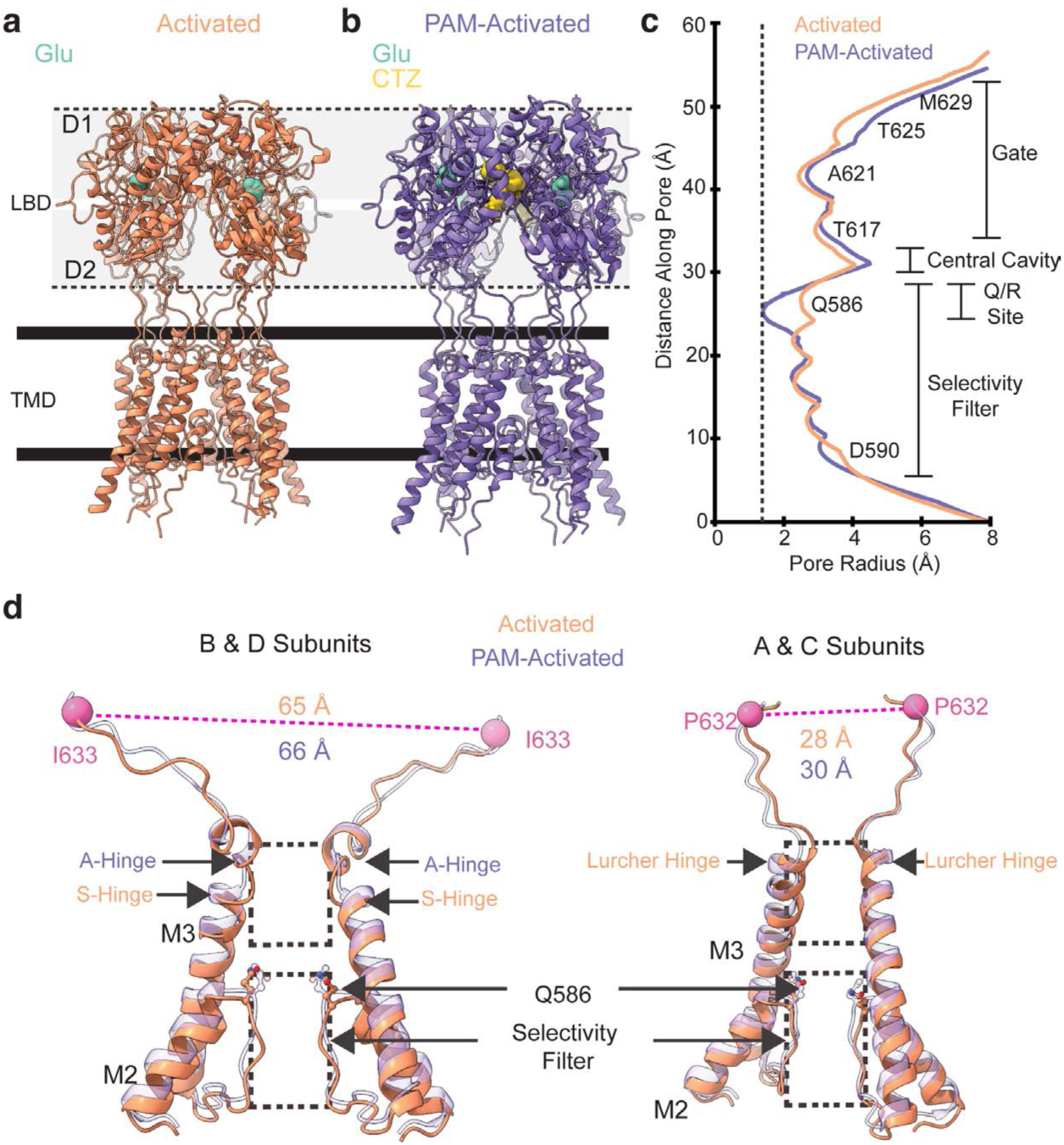
Activated state compared to a PAM-activated state. **a,** Glutamate-activated state of GluA2-γ2 with γ2 excluded. **b,** PAM-activated state of GluA2-γ2 with γ2 excluded (pdb 5WEO). **c**, Pore radius profile comparison between the activated and PAM-activated states. Dashed line represents the 1.4 Å radius of a water molecule. **d**, Comparison between the ion channel pore models of the activated and PAM-activated states. B/D subunits, left; A/C subunits, right.

The ion channel gate is largely similar between the two states (Fig. 4c). The principal difference is at the Q/R site, where the PAM-activated state is significantly more constricted to 1.4 Å versus 2.5 Å in the activated state (Fig. 4c).

In the PAM-activated state, the M3 helices hinge in the B/D subunits at A618, the fourth residue in the SYTANLAAF motif (Fig. 4d). We call this the A-hinge. There is no hinging in the A/C subunits in the PAM-activated state, where in the activated state there is a hinge at the Lurcher site (Fig. 4d). This causes an overall shift in the M3 helices (Fig. 4d) that is likely communicated to M2 and the Q/R site via direct connection, thus reflecting the difference in pore radius at the Q/R site.

The landmark residues in the M3-S2 linker (e.g., I633 & P632) that help facilitate activation (Fig. 3c) have similar degrees of separation in each state, with the PAM-activated state having marginally greater degrees of separation between I633 in B/D (1 Å) and between P632 in A/C (2 Å; Fig. 4d).

The A-hinge in the B/D subunits has been a common feature of all PAM-activated studies that use CTZ^3,6–10,14^. Why is there a difference in pore gating between the states? It is well documented that CTZ enhances activation by preventing desensitization^56^. It is possible that the PAM-activated states captured by cryo-EM represent a post-activated state where the helices have wound back into a relaxed state where the channel is prevented from the conformational transitions that would facilitate desensitization. This idea is consistent with early studies suggesting that CTZ does not contribute to activation of AMPARs, but prevents the desensitized state transition^56–58^. A second possibility is that the glutamate activated state captured here is a higher conductance state (O3 or O4) than the PAM-activated state, and kinking at the S- and Lurcher hinges is required for occupying the high conductance states. This would be consistent with the PAM-activated states occupying O1 or O2 sub-conductance levels^6^.

### Complete reconstruction of AMPAR gating

Isolation of all major gating states from temperature-resolved cryo-EM enables us to completely reconstruct AMPAR gating in the context of just glutamate for the first time. To reconstruct the gating states, in addition to the activated state, we focus on resting-1 and desensitized-2, owing to the data quality over the resting-2 and desensitized-1 counterparts (Extended Data Figs. 4,6). Furthermore, the desensitized-1 and -2 states, as well as resting-1 and -2 states, are largely similar, with RMSD of 0.34 Å and 0.71 Å respectively (Extended Data Fig 9a,b). We focus on the core AMPAR in the center of the GluA2-γ2 complex to discern the gating mechanism (Fig. 5a). Each state maintains a two-fold symmetric tetrameric arrangement.

**Fig. 5.**
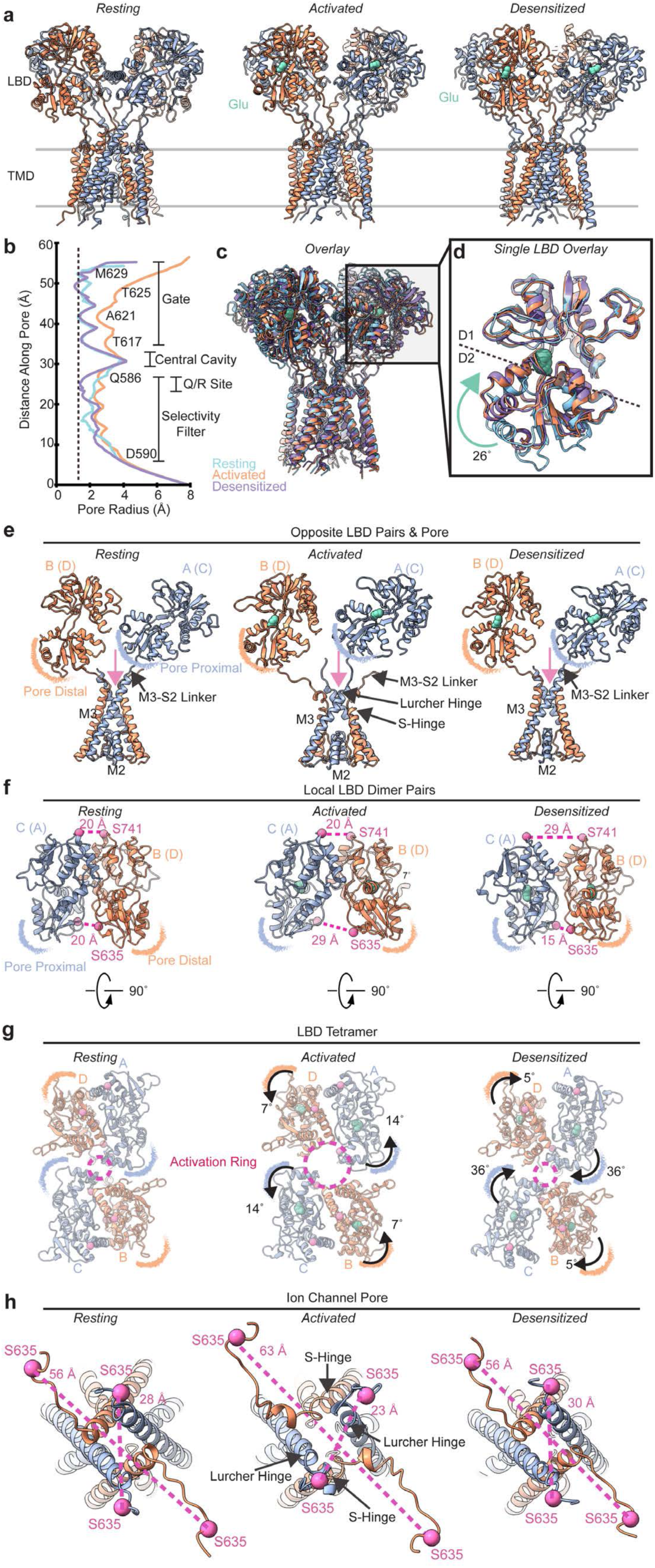
Structural mechanism of glutamate gating. **a,** structures of the resting, activated, and desensitized states from temperature-resolved cryo-EM in this study. γ2 is omitted for clarity and does not undergo observable changes during gating. **b**, pore radius profiles in each gating state. Dashed line represents the 1.4 Å radius of a water molecule. **c**, overlay of each state in panel a, aligned via the TMD. **d**, alignment of a single LBD from each state defines the conformational change associated with glutamate binding. **e**, Opposite LBD pairs (B & A; D & C omitted for clarity) with the ion channel pore below (all subunits) to show the nonequivalent LBD subunit positions. **f**, local LBD dimer pairs with their D1-D1 and D2-D2 distance marker residues in each gating state. **g**, LBD tetramers viewed from the top (90° rotated from panel f) in each gating state. The activation ring between the LBDs dilates during activation. **h**, top view of the ion channel immediately below the LBD tetramer in each gating state. The D2 residue S635 is shown to illustrate the relative positioning of each LBD D2 relative to the central ion channel pore.

The resting (apo) state of the receptor reconstructed here at 42 °C is near identical to the resting state GluA2-γ2 incubated at 4 °C prior to vitrification, where the RMSD between the structures is 1.0 Å (Extended Data Fig. 9c). This is consistent with our temperature steps in single channel currents, where glutamate is the main driver of gating, but temperature augments gating transitions (Fig. 1).

The principal differences in the ion channel pore between the gating states are at the Q/R site and the ion channel gate (Fig. 5b). The map quality in each state enables definitive identification of channel features (Extended Data Figs. 4,6). There are multiple points of channel constriction around the gate in the resting and desensitized states of less than 1.4 Å. While the Q/R site is similar between the resting and activated states, this constricts upon desensitization, perhaps creating a second gate to conductance (Fig. 5b). This is similar to NMDA-subtype iGluRs (NMDARs) studied in the absence of PAMs, where a two gate model was proposed for NMDARs – one gate at the top of NMDAR M3 helices, and the second at the Q/R site equivalent in NMDARs^59,60^.

Overlay of each state highlights the gross differences in the LBD that drive gating of the central ion channel, while the peripheral TM helices are largely similar (Fig. 5c). At the level of the single LBD, both activated and desensitized states show a 26° upward swing of D2 to D1 relative to the resting state, which is unbound to glutamate (Fig. 5d).

Despite the symmetric arrangement, the subunit positions are not equivalent in gating of the channel because of the proximity of the LBDs to the pore (Fig. 5e) coupled with the clamshell closure during glutamate binding (Fig. 5d). When the LBD D2 swings upward to D1 during glutamate binding, this occurs near the channel pore in the A/C subunits. However, in the B/D subunits, this is distal from the pore. Thus, we consider the A/C subunit LBDs pore proximal, and the B/D subunit LBDs pore distal; as the conformational changes coupled with glutamate gating occur, the motions of the B/D subunits apply greater mechanical torque directly to M3 helices via the M3-S2 linkers (below). We hypothesize this is why the B/D subunits cause unwinding further into the SYTANLAAF motif (S-hinge) versus the A/C subunits at the Lurcher Hinge (Fig. 5e).

While the influence of gating is non-equivalent between dimer pairs, coordinated motion of local dimer pairs is critical to the gating process. At the level of local LBD dimers, there are pairs between the B & C subunits and D & A subunits (Fig. 5f). The relative separation between D2s in the local dimer (measured by S635-S635 distances) versus D1s (measured by S741-S741 distances) is how the tension is applied to the linkers below to open the ion channel. During activation, there is a 9 Å increase in the D2 separation in local dimers, which places tension on the LBD-TMD linkers to open the ion channel. In desensitization, the D2-D2 separation is dramatically minimized to 15 Å via maximization D1-D1 separation by 9 Å in local dimers.

Coordination motion of local dimers at the tetrameric level dilates the activation ring between LBDs, which enables torque to be applied to the M3 helices below during gating (Fig. 5g). During activation, the B/D subunits rotate 7° counterclockwise around the gating ring, and the A/C subunits rotate 14 °C counterclockwise relative to the resting state. This collective motion dilates the activation ring. The activation ring is compressed in desensitization by 5° clockwise rotation of the B/D subunits, and 36° clockwise rotation of the A/C subunits. This enables relaxation of the channel below back to an inactivated state despite the LBDs being glutamate bound.

The pore-distal and pore-proximal movements of the LBDs have disparate effects on gating. The collective motions of the local LBDs (Fig. 5f) along the tetrameric LBD layer (Fig. 5g) cause the pore-distal B/D LBDs separate to 63 Å during activation (S635 B to S635 D), which causes unwinding down the M3 helices to the beginning of the SYTANLAAF motif at S615 (Fig. 5h). Distance-wise, the pore proximal LBDs get 5 Å closer (S635 A to S635 C). However, the torque applied by the conformational change of the LBDs causes unwinding to the Lurcher hinge. As the LBDs rotate back during desensitization (Fig. 5g), the helices are wound back, and the gate is restored. Like the hinging of all four M3 helices causing pore dilation at the Q/R site, desensitization appears to narrow the Q/R site to less than 1.4 Å (Fig. 5b). Given the dramatic changes observed at the gate and Q/R site observed here, it is possible that changes at both sites contribute to the sub-conductance observed in iGluRs^6^.

## Discussion

AMPARs, when binding glutamate, transition to other conformational states through a pre-active intermediate^2^ (Fig. 6). This is short-lived, and not captured in this study. It is, however, well-described in others^13,15,61^. Activation is accommodated by D1-D1 contacts between LBDs, an action puts tension on the M3-S2 linkers and opens the channel via the S-hinge (B/D subunits) and the Lurcher hinge (A/C subunits). During desensitization, the LBDs remain glutamate bound, but D2-D2 contacts between LBDs enable relaxation of the tension on M3-S2 linkers, enabling closure of the upper gate, and a finer constriction at the Q/R site. Activation in the presence of a PAM, such as CTZ, captures activated channels where the channel is held open by the A-hinge in the B/D subunits. Negative allosteric modulators (NAMs) such as perampanel and GYKI-52466 bind in the ion channel collar and prevent hinging of the M3 helices, pushing the receptors into an inhibited, desensitized-like state^31,62^.

**Fig. 6.**
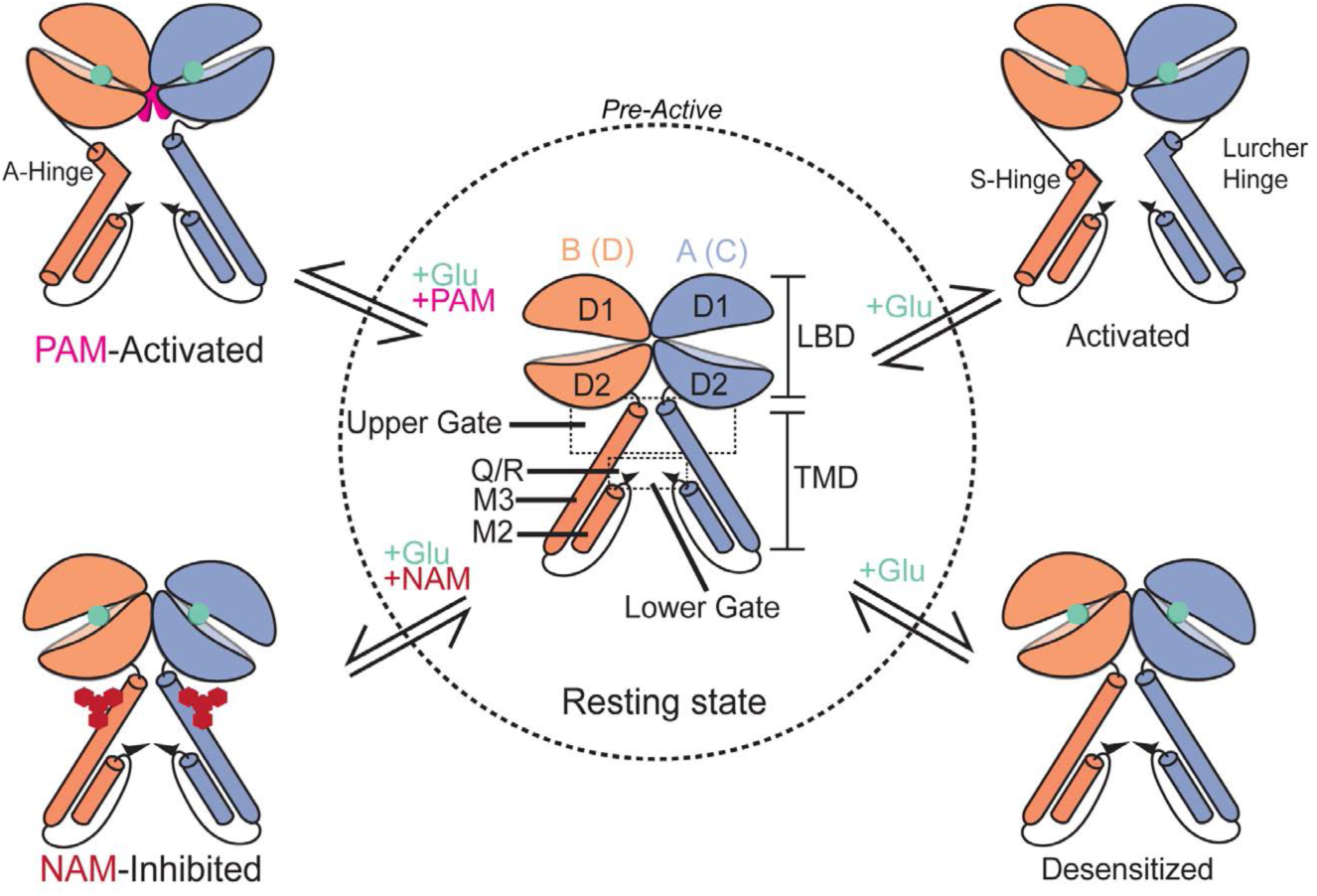
Summary of AMPAR gating and allostery. Transitioning between conformational states of the AMPAR is facilitated by the pre-active intermediate (dashed line). Temperature increases transition probabilities between the resting, activated, and desensitized states. The PAM-activated and NAM-inhibited states are distinct from the normal glutamate gating cycle.

During gating, we see dramatic changes in channel constriction at the Q/R site that point to the site being a lower gate (Fig. 5b), as was suggested for NMDARs^59^. Supporting this idea in AMPARs are recent cryo-EM studies on iGluRs. Kainate-subtype iGluRs (KARs) show dynamic changes at their upper gate as well as their Q/R-site equivalent during gating^12^. Furthermore, in AMPARs structures where the upper gate is open (e.g., ≥ 2.0 Å), only lower conductance levels are observed, potentially because the channel was restricted at the Q/R site^6^. This is highlighted by comparison of the PAM-activated pore to the glutamate activated pore, where there is a 1.1 Å greater constriction at the Q/R site (Fig. 4c). It is possible that dilation of the Q/R site contributes to channel sub-conductance through functioning as a lower gate, and the data here show this possibility.

Our data strongly support that AMPAR gating is temperature sensitive. The augmentation of glutamate responses at physiological temperatures underscores the importance of regulating proper brain temperature, and in considering temperature when studying iGluR function. Our results also show how hyperthermic temperatures dramatically influence AMPAR gating. This hints to how synaptic currents are increased, leading to excitotoxicity, in conditions where hyperthermic temperatures occur in the brain such as traumatic brain injury and many neurological disorders^28,29^.

Using these principles, we use temperature-resolved cryo-EM to capture a glutamate activated state of an iGluR for the first time. For AMPARs, the glutamate activated state is distinct from glutamate activated states in the presence of PAMs. The activated channel hinges in all four pore helices and involves the entirety of the SYTANLAAF motif that is conserved in iGluRs (Extended Data Fig. 8a). Reconstruction of the glutamate gating cycle in AMPARs highlights the nonequivalence of subunit positions, where the B and D subunits unwind the entire SYTANLAAF motif to open the channel, and the A and C subunits unwind to the Lurcher motif. The involvement of the entire SYTANLAAF motif in gating via the S-hinge and hinging at the Lurcher motif in our glutamate activated state is unique from pore opening mechanisms in all iGluR structures, in all which PAMs are utilized (Extended Data Fig. 8c).

It is possible that the GluA2-γ2 activated state we captured with temperature-resolved cryo-EM is distinct from the PAM-activated state because temperature increases channel conductance (Fig. 1), and the PAM-activated states were frozen at lower temperatures for cryo-EM. Indeed, the PAM-activated structures recapitulate lower conductance states O1 and O2^6^. Furthermore, the four-helix hinging we observe in the channel may be distinct conductance state, which would also be supported by the differences in channel constriction at the Q/R site (Fig. 4c), a possible second gate of the channel. This possibility is also likely because most glutamate activated GluA2-γ2 channels occupy O4 in hyperthermic conditions (Table 1).

The recent structure of a PAM-activated NMDAR, where the pore opens via the A-hinge in two subunits (Extended Data Fig. 8c), is constricted at the selectivity filter (Extended Data Fig. 8b). In contrast, the structure of a PAM-activated KAR shows that the pore opens via the L-hinge in the SYTANLAAF motif in all four pore helices (Extended Data Fig. 8c) and has a pore profile like the glutamate activated state captured here (Extended Data Fig. 8b). Thus, we also expect the PAM-activated KAR structure to recapitulate a higher conductance state of the iGluR channel.

The insights from this study will open new avenues for considering temperature in iGluR gating, therapeutic design, and utilizing temperature-resolved cryo-EM to capture states of proteins that are more highly populated at physiological temperatures.

## Methods

### Construct Design

The GluA2-γ2 construct has been well characterized and used in previous studies^3,6,7,30,31,63–65^. In brief, the GluA2* construct, from Yelshanskaya et al. 2014^61^ was fused to the NT of mouse TARPγ2 (NP_031609, CT truncated after L207/TM4). There is a single Gly-Thr spacer between the CT of GluA2* and NT of TARPγ2. After TARPγ2 there is a Thr-Gly-Gly spacer, thrombin cleavage site, enhanced green fluorescent protein (eGFP) for FSEC and monitoring expression, StrepTagII, and stop codon. GluA2-γ2 is in the pEG BacMam vector for baculovirus-driven protein expression in mammalian cells^66^.

### Electrophysiology

HEK 293T cells were plated in poly-D-lysine–coated 35 mm dishes and transfected with GluA2-γ2. Recordings were performed in the outside-out patch-clamp configuration. Pipettes with 8–15 MΩ resistance were filled with 135 mM CsF, 33 mM CsOH,2 mM MgCl2, 1 mM CaCl2, 11 mM EGTA, 10 mM HEPES. The external solution was as follows: 150 mM NaCl, 4 mM KCl, 2 mM CaCl2, 10 mM HEPES, and 10 mM glutamate, pH 7.4. Data were acquired at 50 kHz and low–pass filtered at 5–10 kHz (Axon 200B and Digidata 1550A; Molecular Devices). Pipette holding potential was -100 mV. Data were further filtered at 1 kHz^33^. All recordings were idealized using the segmental k-means algorithm of QuB^67,68^. The kinetic model used three closed and two open levels. After the idealized recording was visually inspected and noise spikes were removed, open and shut times were exported to the Channel Lab program (Synaptosoft), and histograms of the dwell times were displayed and fitted with log-likelihood log-binned subroutines^34^. The mean open time, mean shut time and open probability were obtained using Channel Lab with an imposed dead time of 100 µs. Temperature control was achieved with a micro heating VAHEAT stage (Interherence GmbH, Germany), in addition to a glutamate solution maintained in water baths at each temperature step. To verify temperature, the temperature near the patch was recorded with a BAT-12 Microprobe Thermometer (Physitemp Instruments, LLC) within 5 mm of the microelectrode tip. Details can be found in Table 1.

### Protein Expression and Purification

The GluA2-γ2 bacmid was generated following the method described previously^31^. Following that, ExpiSf9 Cells (Gibco, A35243) cultured at 27 °C, were transfected with the bacmid using ExpiFectamine Sf transfection reagent (Gibco, A38915) to generate the P1 baculovirus. After 5 days, the culture supernatant of ExpiSf9 cells containing the P1 baculovirus was harvested.

The P1 baculovirus supernatant was added to the Expi293F GnTI^-^ cells (Gibco, A39240), maintained at 37 °C in 5% CO_2_ in a 1:10 volume by volume ratio to commence the protein expression. After 16-18 h, the protein expression was induced by adding 10 mM sodium butyrate (Sigma-Aldrich, 303410) and transferring the cells to an incubator set at 30 °C, 5% CO_2_. At this point, 2 µM ZK 20075 (Tocris, 2345) was also added to the cells. After 72 h of infection, the cells were harvested by low-speed centrifugation (∼5,000*g*, 20 mins, 4°C). Cells were washed with 1x PBS (pH 7.4), pelleted again (∼5,000*g*, 20 mins, 4°C), and stored at -80 °C for further use.

For purification, the cell pellet was thawed by rotating in a Tris buffer solution (Tris 20 mM, NaCl 150 mM, pH 8.0) along with the protease inhibitors (0.8 µM aprotinin, 2 µg ml^−1^ leupeptin, 2 µM pepstatin A and 1 mM phenylmethylsulfonyl fluoride). The cells were lysed using a blunt probe sonicator (Fisher Scientific) (3-4 cycles of 1s on, 1s off, ∼20 W power). Lysate was then clarified by low-speed centrifugation (∼2,500*g*, 20 mins, 4 °C) and the membranes were pelleted by using ultracentrifugation (125,000*g*, 45 mins, 4 °C). The pelleted membranes were resuspended, mechanically homogenized and solubilized in the solubilization buffer (150 mM NaCl, 20 mM Tris pH 8.0, 1% *n*-dodecyl-β-D-maltopyranoside (DDM; Anatrace, D310) and 0.2% cholesteryl hemisuccinate Tris salt (Anatrace, CH210) for 2 h at 4 °C with constant stirring. The insoluble material in the solubilized membranes was separated by ultracentrifugation (125,000*g*, 45 mins, 4°C) and the soluble fraction was incubated with the Strep-Tactin XT 4Flow resin (IBA, 2-5010) at 4 °C, rotating overnight. The following day, the protein-bound resin was collected by gravity flow and washed with the GDN buffer containing 20 mM Tris, 150 mM NaCl and 100 µM GDN (Anatrace, GDN101). The protein was eluted with the GDN buffer containing 50 mM Biotin (ThermoFisher, 29129), directly collected in a 100 kDa molecular weight cut-off filter/centrifugal concentrator and concentrated up to ∼500 µl in volume. The eGFP and StrepTag II were cleaved proteolytically, by incubating the protein with thrombin in a ratio of 1:100 (w/w) for 1 h at 25 °C. The sample was then subjected to size-exclusion chromatography by using a Superose 6 increase 10/300 column (Cytiva, 29091596) equilibrated with the GDN buffer. The peak fractions were pooled, concentrated up to ∼4 mg/ml, and used for cryo-EM specimen preparation.

### FSEC

Purified GluA2-γ2 was incubated at three different temperature conditions (4°C, at 37 °C and at 42 °C) for 15 mins, separately and tested for stability by FSEC^69^. FSEC was carried out in an HPLC system attached to a multi-wavelength fluorescence detector, an autosampler (Shimadzu, SIL40C) and a Superose 6 Increase 10/300 GL SEC column (Cytiva). The tryptophan fluorescence was used to monitor the protein stability and retention time (excitation at 280 nm and emission at 325 nm).

### Cryo-EM sample preparation and data collection

Two types of grids were used. C-flat (CF-1.2/1.3-2Au-50, Cat. # CF213-50-Au, Electron Microscopy Sciences) grids were coated with 50 nm Au by Sputter Coater Leica EM ACE600 and plasma cleaned with Ar/O_2_ by Tergeo Plasma Cleaner (Pie Scientific) to make holey 1.2/1.3 gold grids with gold mesh based on published methods^30,70^. These homemade gold grids along with Au-Flat (GF-2/2-2Au-45nm-50, Cat. # AUFT222-50, Electron Microscopy Sciences), were glow discharged in a Pelco Easiglow (25 mA, 120 s glow time and 10 s hold time; Ted Pella, 91000).

The purified GluA2-γ2 protein sample was ultracentrifuged to remove the insoluble material. 20 µl of the protein sample was then incubated on a heat block set at 42 °C for 10 minutes. The protein sample was then immediately spiked with 1 mM glutamate from a stock that was pre-warmed on the 42 °C heat block (for the glutamate-bound sample), and 3 µl of the protein sample was applied to the freshly glow-discharged grids in a FEI Vitrobot Mark IV (Thermo Fisher Scientific) chamber set at 42 °C temperature and 90% humidity and immediately plunge frozen in liquid ethane cooled by liquid nitrogen. For the apo-state GluA2-γ2 condition, the protein was directly applied on grids after incubating on the heat block without spiking in any glutamate.

Data was collected on a Titan Krios G3i microscope (ThermoFisher Scientific) operating at 300kV, equipped with Falcon4i camera and selectris energy filter set at a 10 eV slit width. Data collection was performed in an automated manner using EPU software (ThermoFisher Scientific). A total of 23,236 micrographs were collected for the glutamate spiked condition with a total dose of 40.00 e^−^ per Å^2^, a dose rate of 8.56 e^−^/pixel/s, and a pixel size of 0.97 Å/pixel. For the apo-state, 10,220 micrographs were collected with a total dose of 40.00 e^−^ per Å^2^, a dose rate of 9.92 e^−^/pixel/s, and a pixel size of 0.97 Å/pixel. The defocus range was -1.0 µm to -2.5 µm for all the collections.

### Image Processing

CryoSPARC^71^ (version 4.5.3) was primarily used for all aspects of image processing. Final particle picking was performed with TOPAZ^72^. Details can be found in Extended Data Figures 3-6, Table 2.

### Model Building, Refinements, and Structural Analysis

ChimeraX^73^, ISOLDE^74^, Coot^75^, and PHENIX^76^ compiled by the SBgrid Consortium^77^ were used in combination to perform the model building, refinements, and structural analysis. All visualizations and measurements were performed in ChimeraX. Model quality was assessed with MolProbity^78^. Pore measurements were performed with HOLE^79^. Model quality is reported in Table 2.

### Multiple Sequence Alignment

Rat iGluR protein sequences were accessed from UniProt: (Gria1, P19490; Gria2, P19491; Gria3, P19492; Gria4, P19493; Grik1, P22756, Grik2, P42260; Grik3, P42264; Grik4, Q01812; Grik5, Q63273, Grid1, Q62640; Grid2, Q63226; Nmdz1, P35439; Nmde1, Q00959; Nmde2, Q00960; Nmde3, Q00961; Nmde4, Q62645; Nmd3a, Q9R1M7, Nmd3b, Q8VHN2). Amino acid sequence alignments were generated using Clustal Omega^80^ and were visualized with ESPript 3.0^81^.

## Ethics Declarations

The authors claim no competing interests.

## Data Availability

All cryo-EM reconstructions are deposited into the Electron Microscopy Data Bank (EMDB) and will be released upon publication. The LBD-TMD maps are the primary cryo-EM maps in each deposition and each TMD local map, as applicable, and half maps are supplied as supplemental files in each deposition. All protein models are deposited in the protein data bank (pdb) and will be released upon publication.

## Author Contributions

E.C.T. conceptualized and supervised the project. A.K.M., E.C., V.J., and E.C.T. designed the experiments. A.K.M. performed protein expression, purification, and specimen preparation for cryo-EM. A.K.M collected the cryo-EM data. A.K.M. and E.C.T. processed the cryo-EM data. E.C.T. built the molecular models. E.C. and V.J. designed the electrophysiology experiments. E.C. performed the electrophysiology experiments. E.C. and V.J. performed the electrophysiology data analysis. A.K.M. and E.C.T. wrote the manuscript, which was then edited by all authors.

## Acknowledgements

We thank A. Lau (JHU) and Z. Qiu (JHU) for comments on the manuscript, and W.D. Hale, A. Montaño Romero, and L. Dillard (Twomey Lab) for critical insights during the development of this work. We thank J. F. Cordero-Morales (UTHSC) for lending the VAHEAT micro heating stage used in this work. All cryo-EM data was collected at the Beckman Center for Cryo-EM at Johns Hopkins. E.C.T. is supported by National Institutes of Health (NIH) grant R35GM154904, the Searle Scholars Program (Kinship Foundation #22098168) and the Diana Helis Henry Medical Research Foundation (#142548). V.J. is supported by NIH grant R35GM122528.

## Extended Data Figures

**Extended Data Fig. 1.**
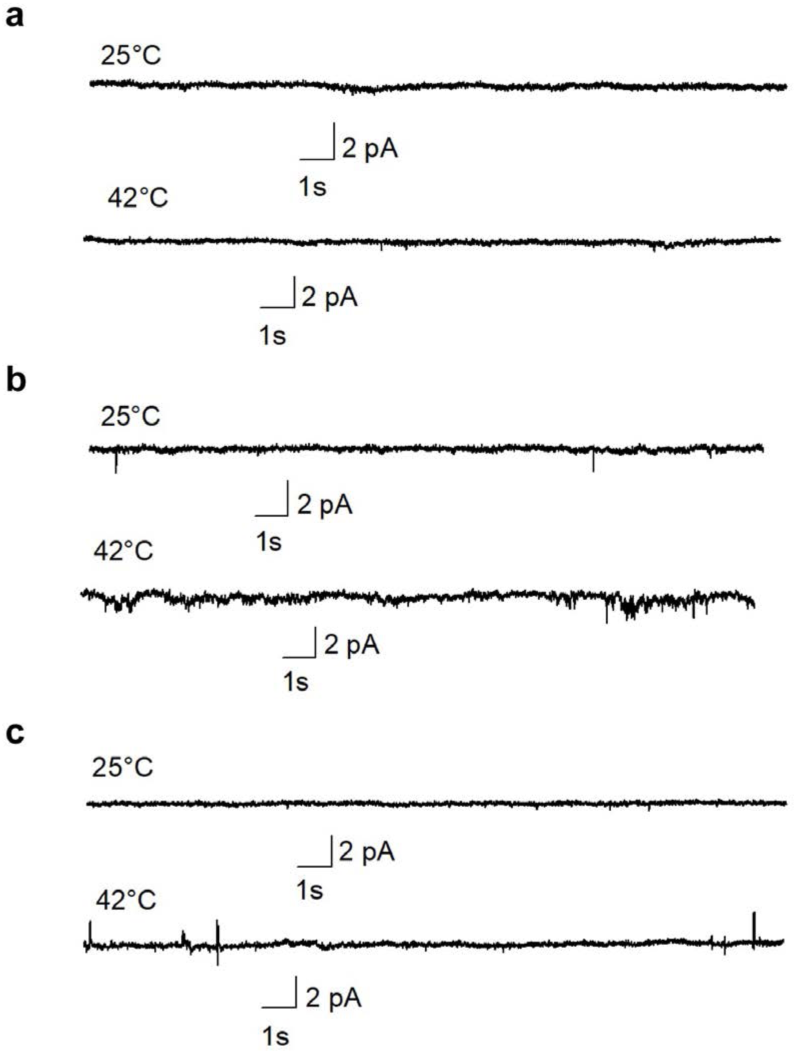
Control current recordings from untransfected cells. **a**,**b,c**, Three separate patches where currents were recorded at 25°C and 42°C in the presence of 10 mM glutamate from HEK293T cells not transfected with GluA2-γ2.

**Extended Data Fig. 2.**
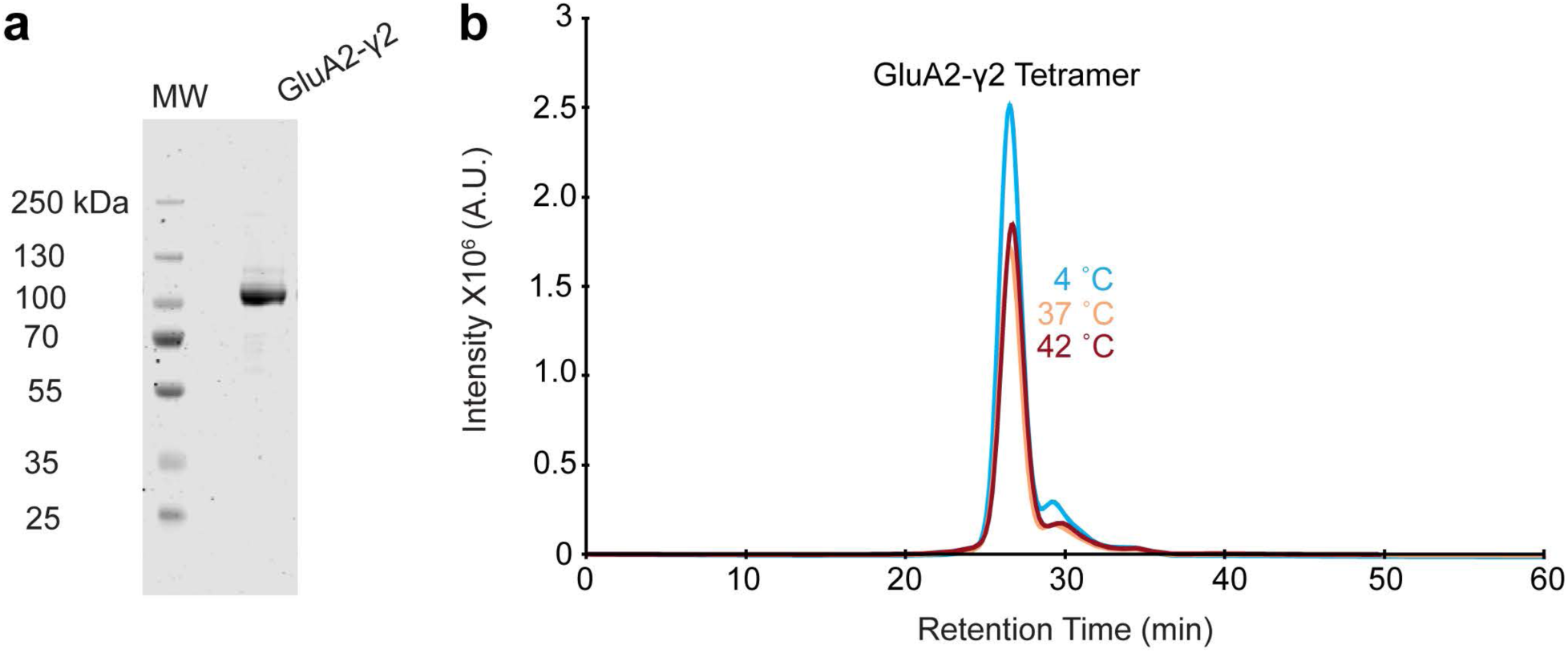
Purification and stability of GluA2-γ2 at physiological temperatures. **a**, SDS-PAGE of purified GluA2-γ2. Molecular weight marker (left), GluA2-γ2 (right). **b**, FSEC tryptophan fluorescence chromatograms of purified GluA2-γ2 at 4 °C, 37 °C, and 42 °C.

**Extended Data Fig. 3.**
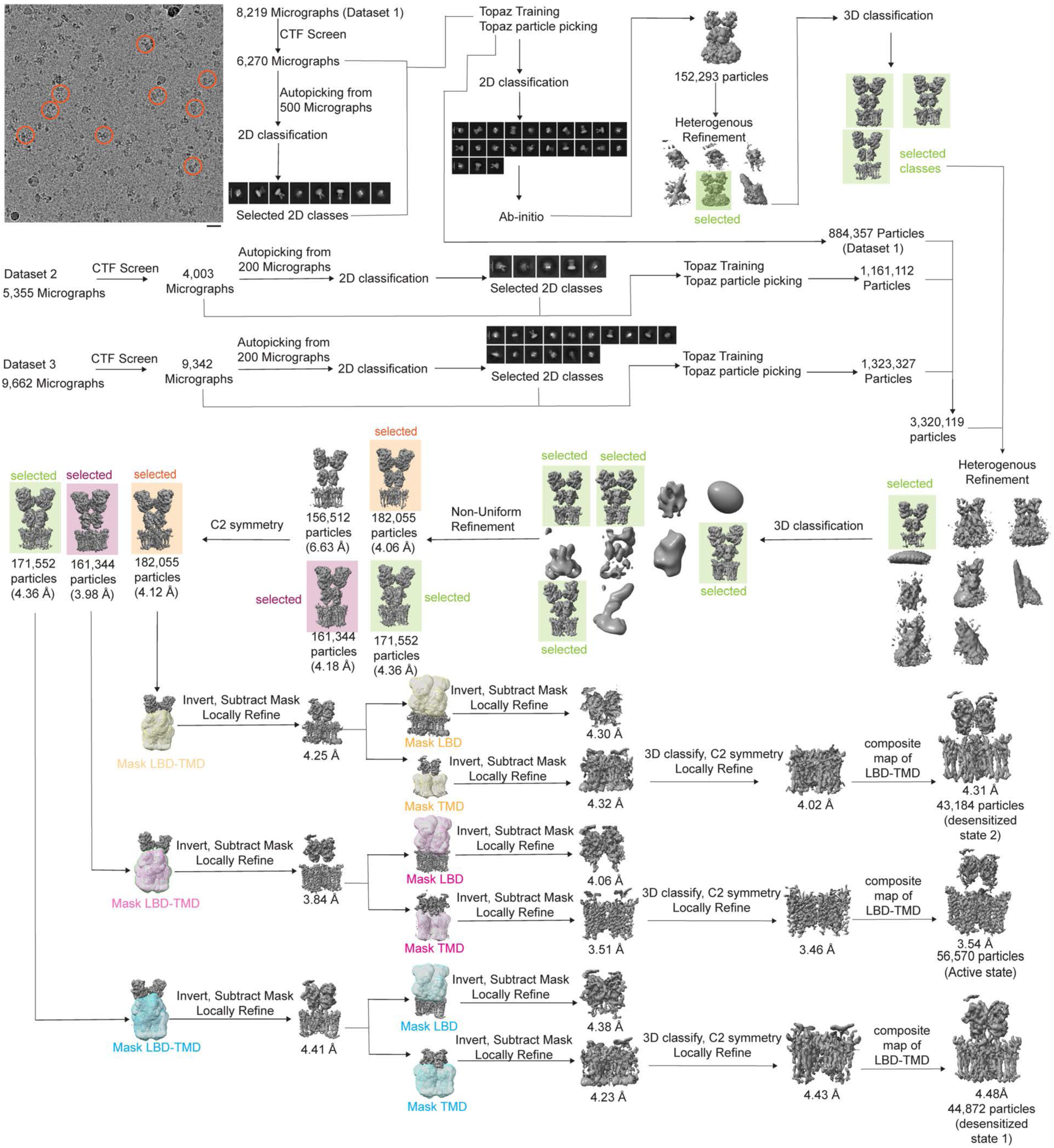
Cryo-EM image processing workflow for Glu-42°C data.

**Extended Data Fig. 4.**
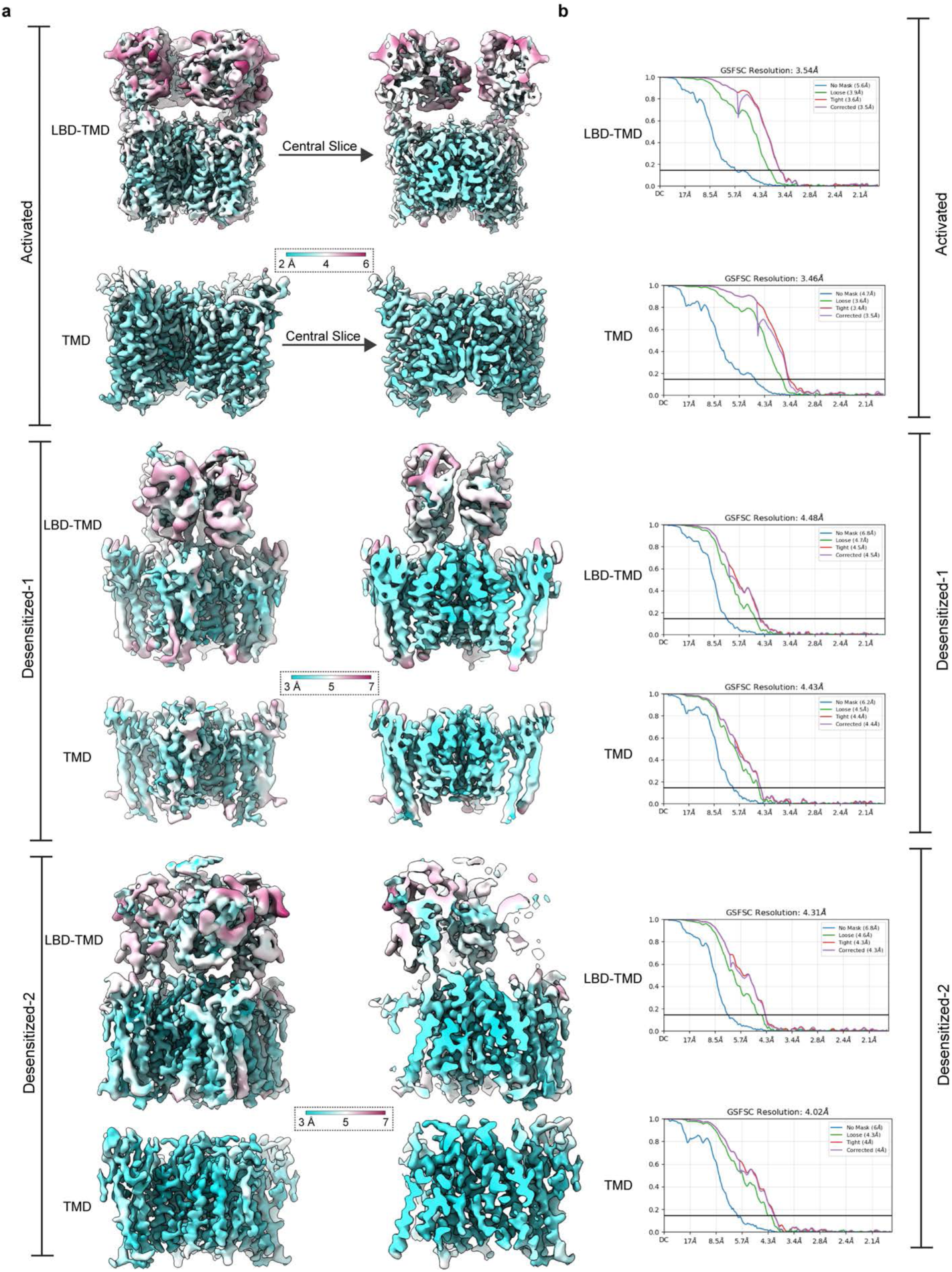
Local and overall map qualities for glutamate 42°C cryo-EM data. **a**, local resolution maps computed for each voxel in activated, desensitized-1, and desensitized 2 maps, where resolutions were computed at Fourier-shell correlation (FSC)=0.143. **b**, Overall cryo-EM map (LBD-TMD, TMD) FSC plots. Black line is FSC=0.143.

**Extended Data Fig. 5.**
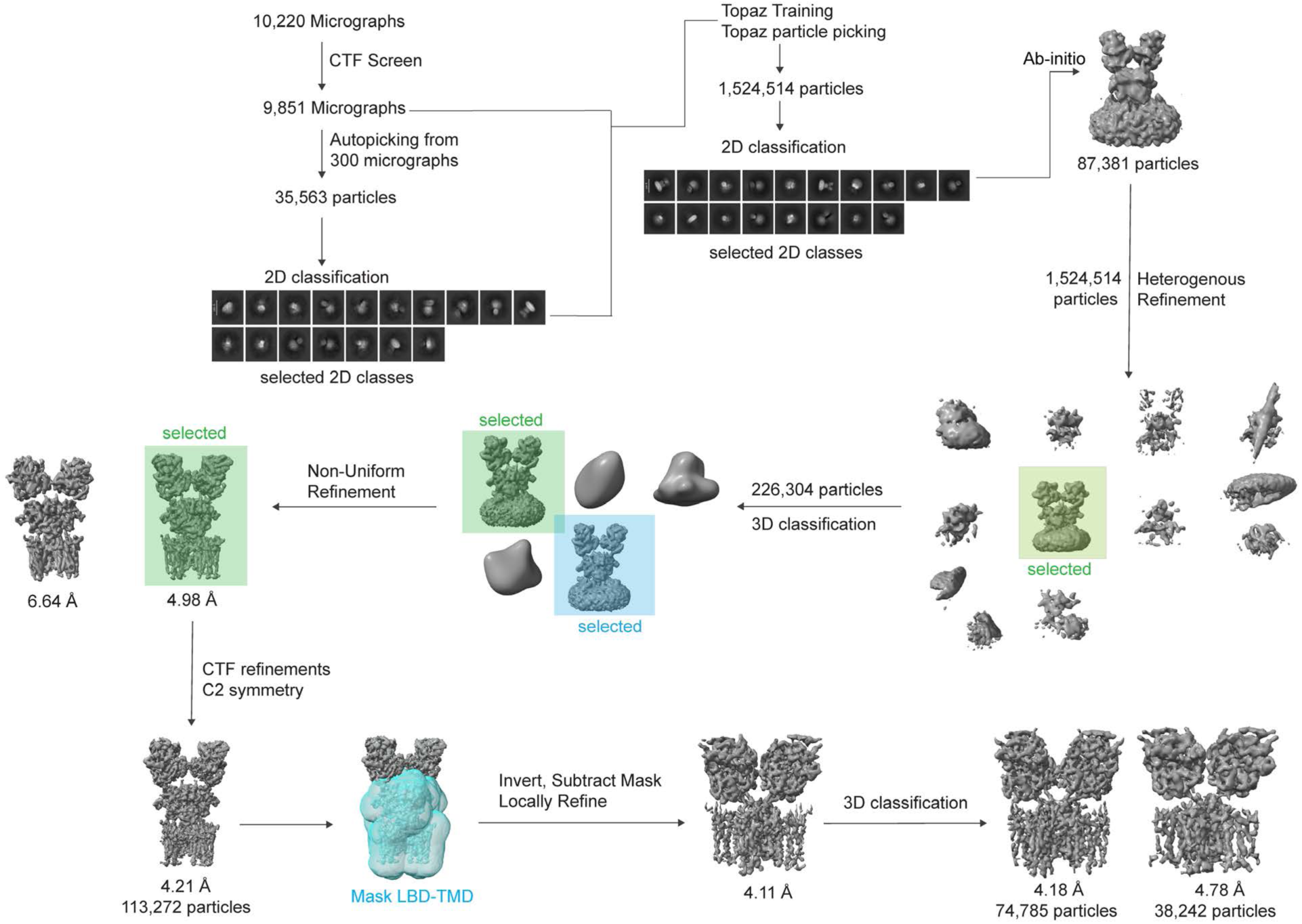
Cryo-EM image processing workflow for resting state 42°C data.

**Extended Data Fig. 6.**
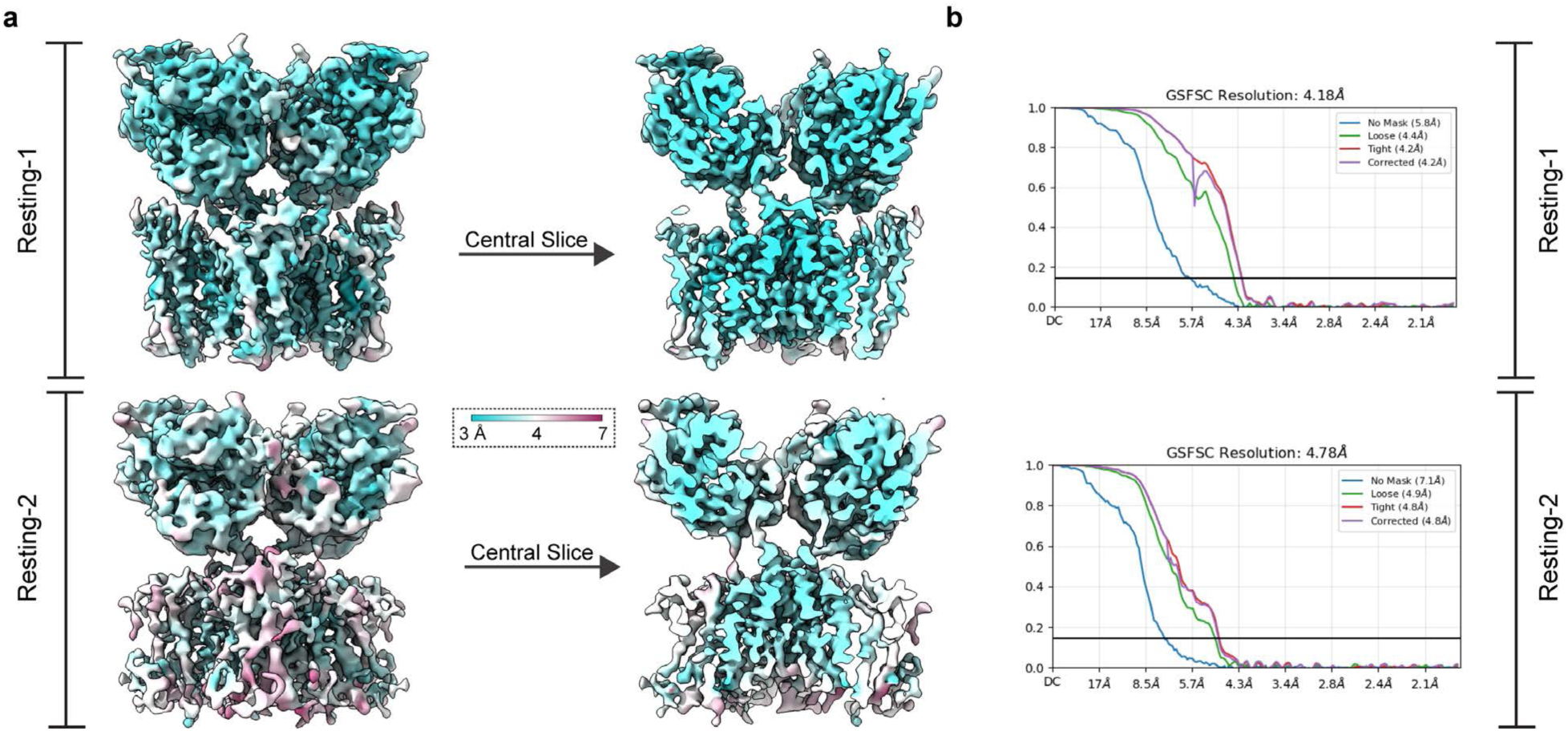
Local and overall map qualities for resting state 42°C cryo-EM data. **a**, local resolution maps computed for each voxel in resting-1 and -2 maps, where resolutions were computed at FSC=0.143. **b**, Overall cryo-EM map (LBD-TMD, TMD) FSC plots. Black line is FSC=0.143.

**Extended Data Fig. 7.**
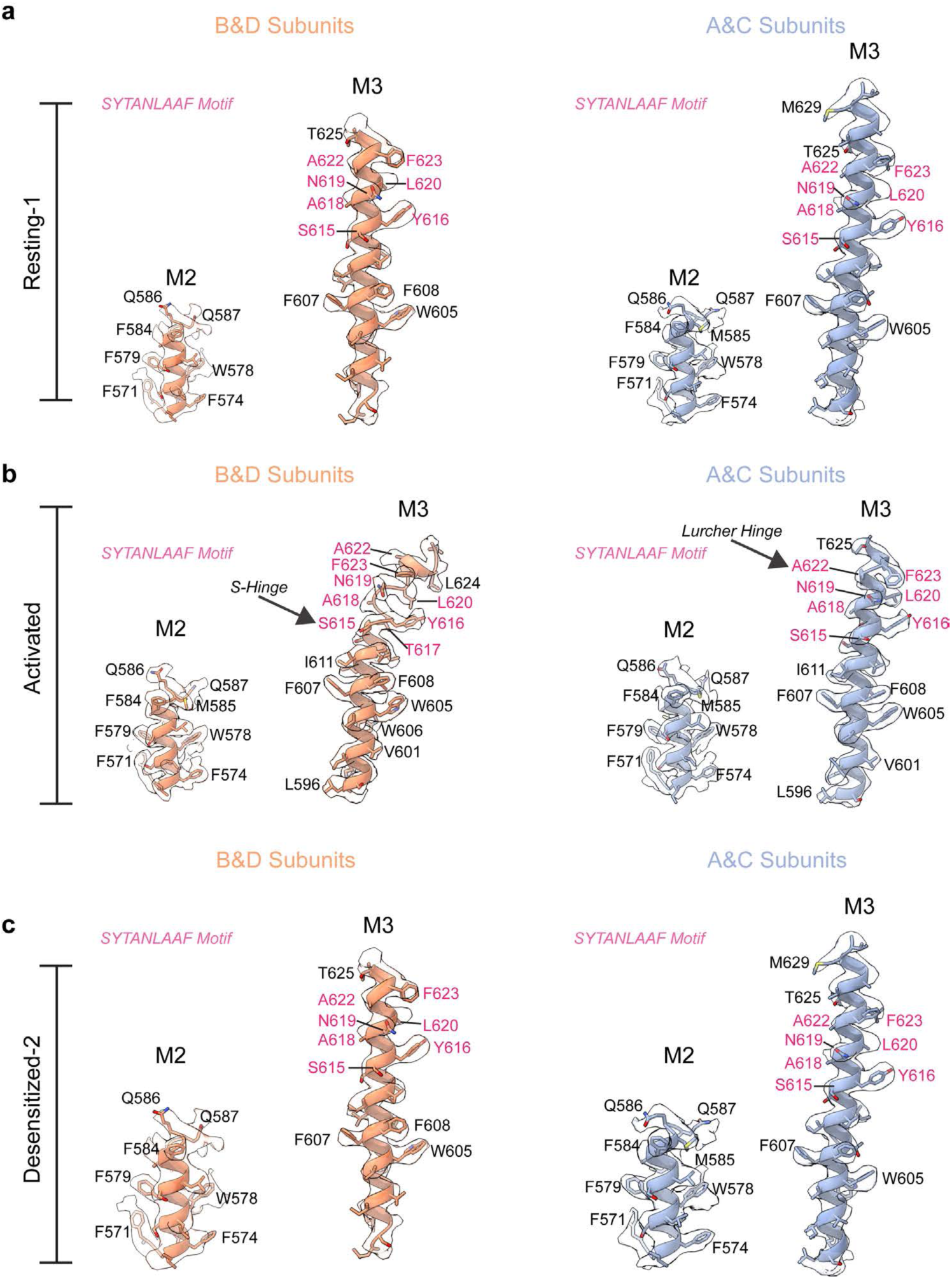
Pore helix map quality and model fit. **a,** Resting -1 M2 and M3 model-map fit, B&D subunits on left, A&C subunits right. **b**,**c** as in panel a, but for activated and desenstizied-2 states, respectively. SYTANLAAF motif is in pink in all panels.

**Extended Data Fig. 8.**
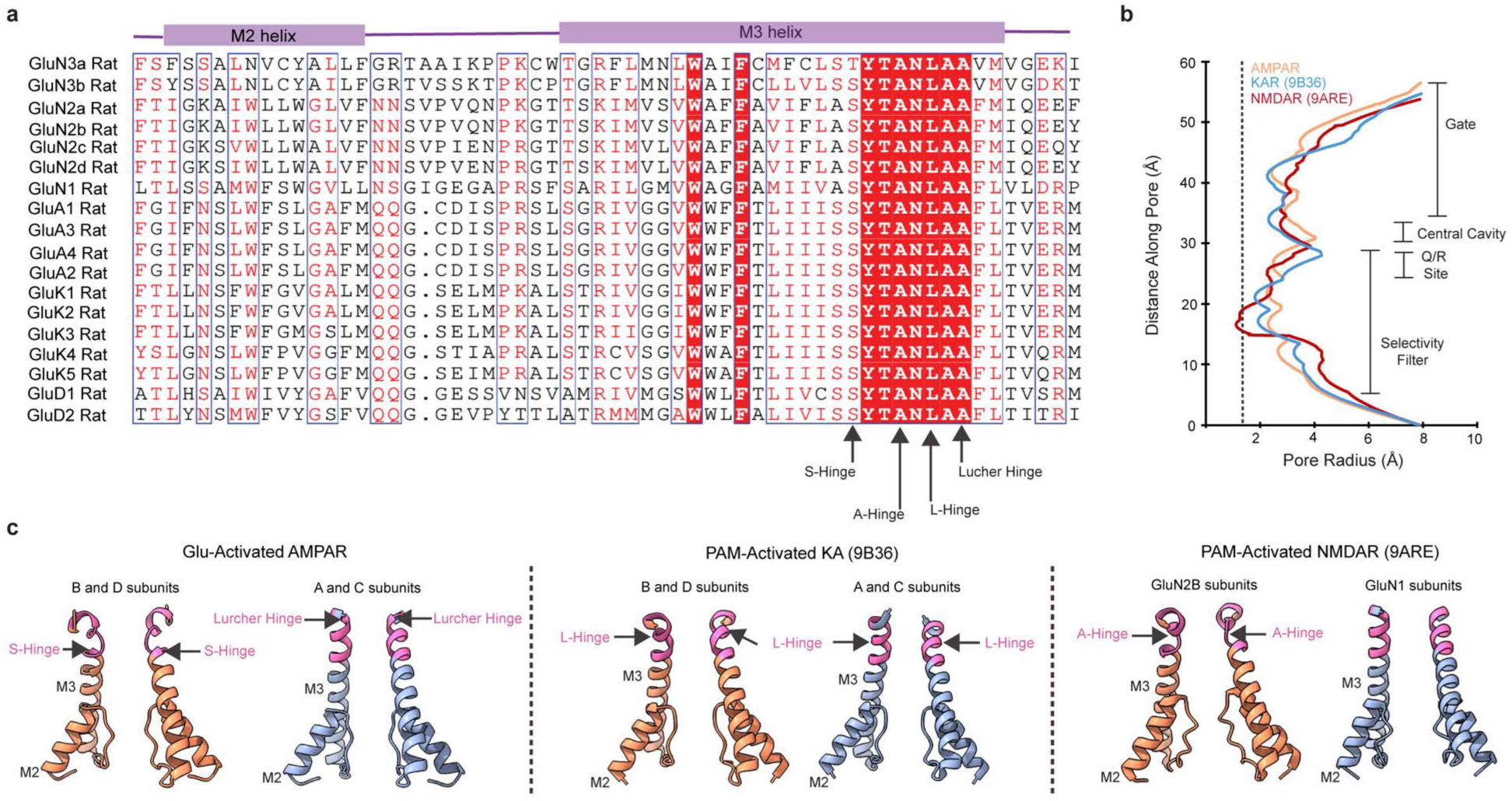
Conserved pore motifs in iGluRs and pore features of activated iGluRs. **a**, Amino acid sequence alignment of major iGluR families. **b**, Pore profiles of PAM-activated KAR (pdb 9B36) and PAM-activated NMDAR (pdb 9ARE) compared to the glutamate-activated state from this study. **c**, Pore hinging in structures plotted in panel B. Hinging locations are also marked in panel a.

**Extended Data Fig. 9.**
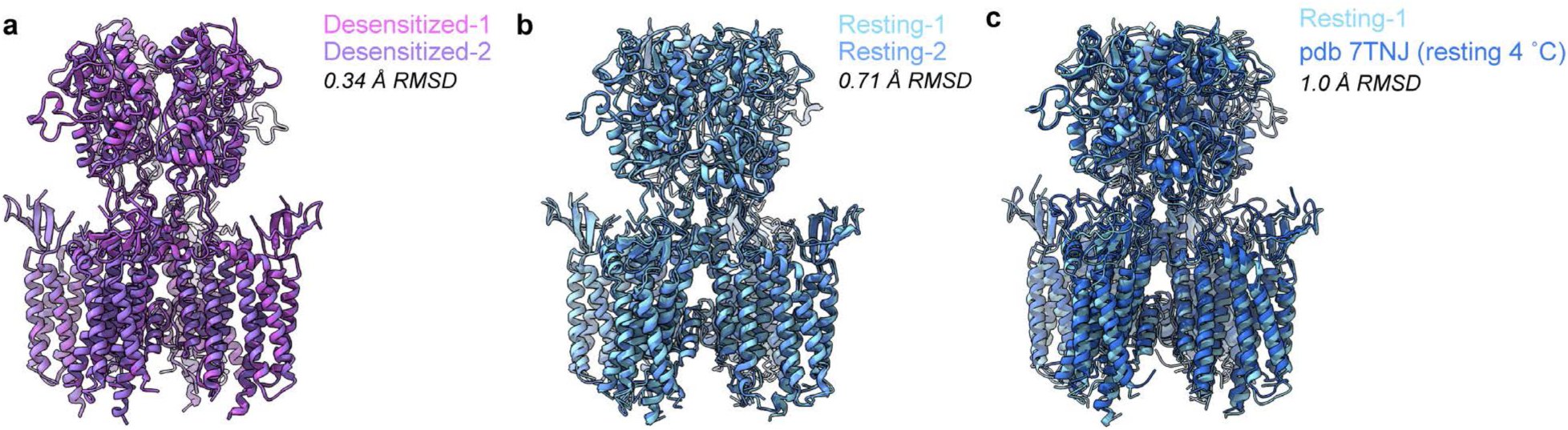
Structural comparison between temperature-resolved cryo-EM states, and comparison between 42 °C and 4 °C GluA2-γ2 resting states. **a**, Structural alignment between Desensitized-1 and -2. **b**, Structural alignment between Resting-1 and -2. **c**, Structural alignment between Resting-1 and GluA2-γ2 resting prepared at 4 °C (pdb 7TNJ).

